# Harnessing machine learning to unravel protein degradation in *Escherichia coli*

**DOI:** 10.1101/2020.10.04.325795

**Authors:** Natan Nagar, Noa Ecker, Gil Loewenthal, Oren Avram, Daniella Ben-Meir, Dvora Biran, Eliora Ron, Tal Pupko

## Abstract

Degradation of intracellular proteins in Gram-negative bacteria regulates various cellular processes and serves as a quality control mechanism by eliminating damaged proteins. To understand what causes the proteolytic machinery of the cell to degrade some proteins while sparing others, we employed a quantitative pulsed-SILAC (Stable Isotope Labeling with Amino acids in Cell culture) method followed by mass spectrometry analysis to determine the half-lives for the proteome of exponentially growing *Escherichia coli*, under standard conditions. We developed a likelihood-based statistical test to find actively degraded proteins, and identified dozens of novel proteins that are fast-degrading. Finally, we used structural, physicochemical and protein-protein interaction network descriptors to train a machine-learning classifier to discriminate fast-degrading proteins from the rest of the proteome. Our combined computational-experimental approach provides means for proteomic-based discovery of fast degrading proteins in bacteria and the elucidation of the factors determining protein half-lives and have implications for protein engineering. Moreover, as rapidly degraded proteins may play an important role in pathogenesis, our findings could identify new potential antibacterial drug targets.

## Introduction

The degradation of intra-cellular proteins is a fundamental process of life and serves various important physiological functions, including removal of abnormal proteins and regulation of basic cellular processes (Goldberg, 2003; Rubinsztein, 2006; Gsponer *et al*, 2008; Hosoi & Ozawa, 2010; Gur *et al*, 2011; Maupin-Furlow, 2012; Mahmoud & Chien, 2018). In eukaryotes, the covalent binding of a small protein, ubiquitin, marks proteins for degradation by the proteasome (Hershko, 1991). In bacteria, ATP-dependent AAA+ (ATPases associated with cellular activities) proteases use ATP hydrolysis to fuel substrate degradation (Mahmoud & Chien, 2018). Degradation of intracellular proteins in Gram-negative bacteria is mainly performed by five ATP-dependent AAA+ proteases: ClpAP, ClpXP, Lon, HslUV and FtsH (Baker & Sauer, 2006; Gur *et al*, 2011; Mahmoud & Chien, 2018).

Since protein degradation is an irreversible process with a considerable damaging potential (Conlon *et al*, 2013), protease activity has to be carefully regulated. Many factors were suggested for regulating degradation, mainly for eukaryotic cells. These include physical properties such as protein mass, isoelectric point, surface-accessibility, structural disorder, and low-complexity regions (Dice *et al*, 1979; Miller *et al*, 1987; Tompa *et al*, 2008; van der Lee *et al*, 2014), as well as sequence-related properties such as the N-end rule, PEST (sequence that is rich in proline, glutamic acid, serine, and threonine), destruction-box, KEN-box, and other sequence motifs (Bachmair *et al*, 1986; Rogers *et al*, 1986; Hoskins *et al*, 2000; Ishii *et al*, 2000; Flynn *et al*, 2003; Burton *et al*, 2005; Shah & Wolf, 2006). Sequence motifs that are involved in the regulation of protein degradation are known as “degrons”. It is assumed that these sequences are located at the C- and N-termini of proteolytic substrates (Bachmair *et al*, 1986; Keiler *et al*, 1996; Flynn *et al*, 2003; Koren *et al*, 2018; Lin *et al*, 2018). For example, it was suggested that ClpXP recognizes proteolytic substrates through five degron classes; three are located at the N-terminus of proteins: polar-T/ϕ-ϕ-basic-ϕ, NH_2_-Met-basic-ϕ-ϕ-ϕ-X5-ϕ, ϕ-X-polar-X-polar-X-basic-polar (where ϕ represents hydrophobic amino acids, and X any amino acid), and two are located at the C-terminus: LAA-COOH and RRKKAI-COOH. A proteolytic substrate can bear either a C-terminal motif, an N-terminal motif, or both (Flynn *et al*, 2003). The LAA-COOH motif is similar to the ssrA tag (AANDENYALAA), which is known to be appended to the C-terminus of proteins for which translation cannot be completed (Keiler *et al*, 1996), thereby targeting the tagged, defected protein to degradation by the ClpXP protease (Flynn *et al*, 2001, 2003). Several attempts have been made to systematically estimate the collective and/or individual contribution of known degradation regulating factors. This was done either in the context of the overall variability observed for protein stability in bacteria (Bachmair *et al*, 1986), or in the context of the substrate repertoires of specific proteases (Flynn *et al*, 2003; Burton *et al*, 2005; Gur & Sauer, 2008; Arends *et al*, 2018).

Over the past decade, it has become possible to track protein degradation *in-vivo* at the global level, i.e., degradation profiles (Boisvert *et al*, 2012; Jovanovic *et al*, 2015). These profiles were determined by the heavy – light amino acid pulse SILAC technology followed by quantitative mass spectrometry (MS) (Schwanhäusser *et al*, 2009) as well as other MS-based methods, for various organisms and in different physiological contexts (Flynn *et al*, 2003; Price *et al*, 2010; Schwanhäusser *et al*, 2011; Westphal *et al*, 2012; Michalik *et al*, 2012; Christiano *et al*, 2014; Mathieson *et al*, 2018; Swovick *et al*, 2018; Arends *et al*, 2018). In the pulsed-SILAC setting, the isotopic ratios of the different mass-labels, which are frequently used for differential expression analysis, are instead used to determine the dynamics of protein degradation (Mann, 2006). We used pulsed SILAC to determine protein half-lives in exponentially growing *E. coli*. We then used statistical modeling of protein stability to classify each protein to one of three mutually exclusive stability groups that we termed as stable, slow-degrading and fast-degrading proteins. We next searched for various features that characterize each of these stability groups and used them for training a machine learning classifier.

Machine-learning approaches have been proved useful for predicting various aspects of protein functions, including prediction of novel effector proteins in pathogenic bacteria, prediction of phosphorylation sites, and prediction of subcellular locations, to name but a few (Hayes & Borodovsky, 1998; Burstein *et al*, 2009, 2012; Miller *et al*, 2009; Nanni *et al*, 2012; Teper *et al*, 2016; Cheng *et al*, 2018). A critical requirement for such a machine-learning approach is a reliable training data, which in the context of protein degradation is accurate determination of protein half-lives. To this end, we used our data to develop machine-learning classification algorithms to assign each cellular protein to one of the stability groups, based on its associated features and reached an Area Under the ROC Curve (AUC) of 0.72.

## Materials and Methods

### Reagents and bacteria

MgSO_4_, NaCl, NH_4_Cl, CaCl2, glucose, thiamine and light (Lys0) L-lysine were purchased from Merck (Burlington, MA, USA). Na_2_HPO_4_ · 7H2O and KH_2_PO_4_ were purchased from Thermo Fisher Scientific (Waltham, MA, USA) and Avantor (Radnor, PA, USA). Medium (Lys6) and heavy (Lys8) isotopes were purchased from Cambridge Isotope Laboratories (Tewksbury, MA, USA). *E. coli* K-12 auxotrophic for lysine (strain JW2806-1, from the Keio collection of single gene knockouts) was employed in the experiments conducted in this study.

### Bacterial cell culture and pulsed SILAC labeling

*E. coli* cultures were grown over night on M9 medium (x5 M9 salts (0.24 M Na_2_HPO_4_ · 7H_2_O, 0.11 M KH_2_PO_4_, 42.8 mM NaCl, 93.45 mM NH_4_Cl), 2 mM MgSO_4_, 0.4% glucose, 0.1 mM CaCl_2_, 0.1 mg/ml thiamine) agar plates supplemented with 250 μg/ml lysine and 50 μg/ml kanamycin. For isotope labeling two single colonies were passaged twice at 37°C on M9 medium containing 250 μg/ml SILAC residues, either light (L), or medium (M). Samples from the two cultures were then reseeded at a low optical density (OD_M_600 nm = 0.033, OD_L_600 nm = 0.03) in fresh M or L M9 medium. Upon early-log phase (OD_M_600 nm = 0.343, OD_L_600 nm = 0.267), two samples were taken; one of M labeled cells that was used for verification of full incorporation of the M lysine isotope (>98% incorporation). The second sample was a mixture of equivalent amounts of cells from the L and M cultures (t_0 h_). At that time point, the M-containing culture medium was replaced with an equivalent volume of “heavy” (H)-containing medium (250 μg/ml), while the L-containing culture medium was replaced with an equivalent volume of fresh L-containing medium, using rapid filtration on 0.22 μm filters. Following medium exchange, the culture now growing in H medium was sampled at five time points (t_0.25 h_, t_1 h_, t_2 h_, t_3 h_, t_4 h_) and mixed with an equivalent amount of cells growing in the L medium. The cells were harvested by centrifugation at 4,000 g, 4°C for 10 min, resuspended in 1 ml of M9 medium, snap-frozen in liquid nitrogen, and stored at −80°C. The experimental setup is illustrated in Fig. 1.

**FIG. 1.**
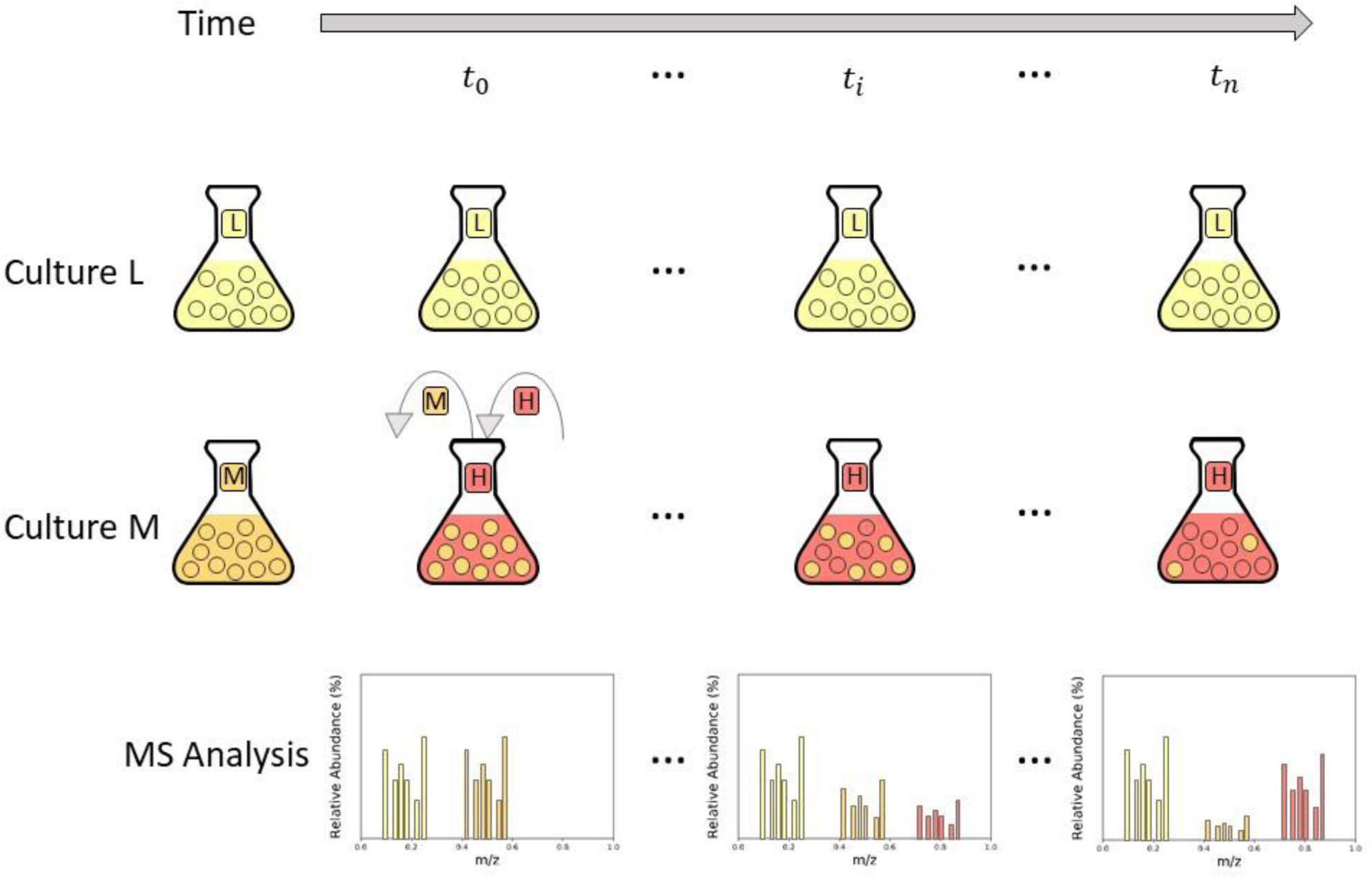
Pulsed-SILAC method illustration. *E. coli* cells are cultured in different SILAC media (Culture L and Culture M) containing either “light” (yellow) or “medium” (orange) lysine until full incorporation of the relevant isotope (leftmost Erlenmeyer flask in each Culture). The grey arrow at the top of the figure represents the experiment timeline (during bacterial exponential-growth phase). At time *t*_0_, the “medium” lysine isotope of Culture M is replaced by “heavy” lysine isotope (red). Next, at each time point *t*_*i*_ (including *t*_0_ and *t*_*n*_), equal amounts of cells are sampled from Culture L and Culture M, mixed, and analyzed by Mass-Spectrometry (MS). The resulting ratios of M/L isotopes over time measures the rate of protein degradation.

### Proteomics

Sample preparation, liquid chromatography, mass spectrometry, and data processing were done at the De Botton Protein Profiling institute of the Nancy and Stephen Grand Israel National Center for Personalized Medicine, Weizmann Institute of Science.

### Sample preparation

All chemicals were purchased from Sigma-Aldrich unless otherwise noted. Cell pellets were lysed with 5% SDS in 50 mM Tris-HCl. Lysates were incubated at 96°C for 5 min, followed by six cycles of 30 s of sonication (Bioruptor Pico, Diagenode, USA). Protein concentration was measured using the BCA assay (Thermo Scientific, USA) and a total of 30 μg protein was reduced with 5 mM dithiothreitol and alkylated with 10 mM iodoacetamide in the dark. Each sample was loaded onto S-Trap microcolumns (Protifi, USA) according to the manufacturer’s instructions. In brief, after loading, samples were washed with 90:10% methanol/50 mM ammonium bicarbonate and digested with LysC (1:50 protease/protein) for 1.5 h at 47°C. The digested peptides were eluted with 50 mM ammonium bicarbonate and incubated overnight with trypsin at 37°C. Two additional elutions were performed using 0.2% formic acid and 0.2% formic acid in 50% acetonitrile. The three elutions were pooled together and vacuum-centrifuged to dry. Samples were kept at −80 °C until analysis.

### Liquid chromatography

ULC/MS grade solvents were used for all the chromatographic steps. Each sample was loaded using split-less nano-Ultra Performance Liquid Chromatography (10 kpsi nanoAcquity; Waters, Milford, MA, USA). The mobile phase was: (A) H_2_O + 0.1% formic acid and (B) acetonitrile + 0.1% formic acid. Desalting of the samples was performed online using a reversed-phase Symmetry C18 trapping column (180 µm internal diameter, 20 mm length, 5 µm particle size; Waters). The peptides were then separated on a T3 high strength silica nano-column (75 µm internal diameter, 250 mm length, 1.8 µm particle size; Waters) at 0.35 µL/min. Peptides were eluted from the column into the mass spectrometer using the following gradient: 4% to 25% buffer B in 155 min, 25% to 90% buffer B in 5 min, maintained at 90% for 5 min and then back to initial conditions.

### Mass spectrometry

The nanoUPLC was coupled online through a nanoESI emitter (10 μm tip; New Objective; Woburn, MA, USA) to a quadrupole orbitrap mass spectrometer (Q Exactive HF, Thermo Scientific) using a FlexIon nanospray apparatus (Proxeon). Data were acquired in data dependent acquisition (DDA) mode, using a Top20 method. MS1 resolution was set to 120,000 (at 400 m/z), mass range of 375-1650 m/z, automatic gain control of 3E6 and maximum injection time was set to 60 msec. MS2 resolution was set to 15,000, quadrupole isolation 1.7 m/z, AGC of 1e5, dynamic exclusion of 45 sec and maximum injection time of 60 msec.

### Data processing

Raw data were processed with MaxQuant version 1.6.0.16 (Cox & Mann, 2008). The data were searched with the Andromeda search engine (Cox *et al*, 2011) against the Uniprot *E. coli* K-12 proteome database (UP000000625) appended with common lab protein contaminants and the following modifications: Carbamidomethylation of C as a fixed modification and oxidation of M and deamidation of N and Q as variable ones. Labeling was defined as H-heavy K8, M – medium K4 and L – light K0. The match between runs option was enabled as well as the re-quantify function. The rest of the parameters were used as default. Decoy hits were filtered out using Perseus version 1.6.0.7 (Tyanova *et al*, 2016), as well as proteins that were identified on the basis of a modified peptide only. The H/L, H/M and M/L ratios and raw intensities in the “proteinGroups” file were used for further calculations (**Supplemental Dataset 1**).

### Determination of protein half-life

We used a modeling scheme similar to that described by Boisvert *et al*. (2012). As stated above, we sampled bacteria from two different cultures, grown in either L- or M-containing media. At time 0, the M-containing medium was replaced with H-containing medium. Let 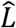 be the abundance of the L isotope in cells grown in L-containing medium (i.e., the number of protein molecules harboring the L isotope). We assume that in each generation, the number of cells is doubled and consequently, 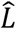 is doubled as well. Let *t*_*cc*_ be the generation time in minutes (∼60 minutes in our cultures). Thus, when the cells are growing for *t* minutes, the number of generations is *t*/*t*_*cc*_ and the total abundance of the integrated L isotope is:

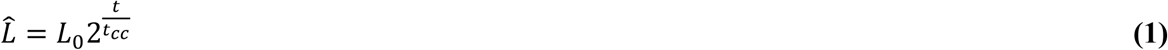

Let 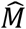 be the abundance of the M isotope in cells grown in M-containing medium. Following removal of the M-containing medium at time 0, 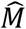 is expected to have an exponential decay with a specific rate factor. We note that cell division does not affect 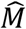, because 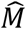 measures the total amount of M in the cells. Thus, 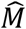 is expected to decrease due to protein degradation according to the following equation:

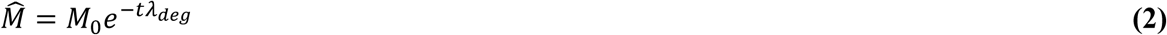

The parameter *λ*_*deg*_ governs the degradation rate. High values of *λ*_*deg*_ indicate higher rates of degradation, and at the limit, when *λ*_*deg*_ = 0, the abundance 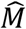 remains M_0_ regardless of *t*.

Up until the medium replacement step 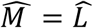 (because these two isotopes are used in parallel under the same conditions). Upon medium replacement, the M isotope available in the medium is washed away by filtration and replaced with H isotope containing medium. We do the same procedure for the L isotope: The L medium is washed away and replaced with fresh L containing medium (**Fig. 1**). Thus, at the replacement time point, 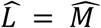 and after this time point, added L in the cells is the same as added H in the cells. Thus, the total abundance of L in the cells should equal the sum of the integrated M and H isotopes:

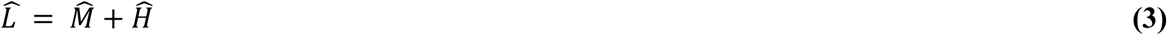

Taking the next samples, at each time point, we made sure to take the same number of cells from the L culture and from the H+M culture. Hence the measured levels of L and M, at time point *t* are:

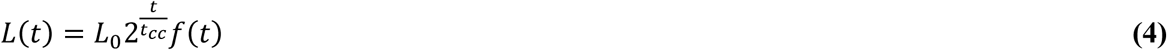

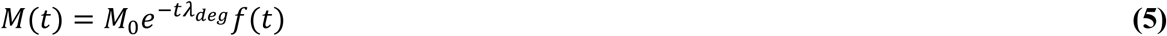

where *f* (*t*)is the fraction of cells sampled at time *t*. From these equations, we obtain:

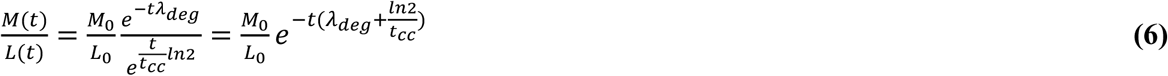

In our experiments, the proteomic results after MaxQuant analysis provide us with the 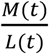 and 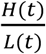 observed values. In theory, according to equation (3) these two ratios should sum up to 1. In practice, however, small deviations from the sum of 1 are observed (0.99 ± 0.01, at 95% confidence interval). Hence, we add a normalization step in which we multiply both ratios by a fixed constant so that they sum to 1, for every *t*. Also note that according to the experimental design, M_0_ should equal L_0_ and thus, their ratio should be 1. In our experiment, we observed a ratio of 1.02 ± 0.02, at 95% confidence interval. The normalized 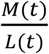 ratios are plotted against *t*, where *M*(*t*) and *L*(*t*) represent the observed intensity of the medium and light isotopes at each time point. Using R’s nonlinear least squares routine, *nls* (Jones *et al*, 2001), we then fit the obtained curve to a simple exponential function of the form:

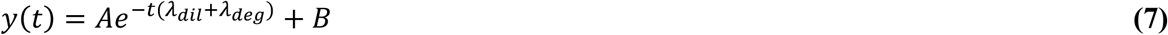

The estimated parameters in this non-linear regression are *A, B* and *λ*_*deg*_. Comparing equations **(6)** and **(7)**, *A* corresponds to the normalized 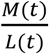 ratio at *t* =0, *λ*_*deg*_ corresponds to the degradation constant and *B*accounts for the offset seen in data, which is attributed to recycling by Boisvert *et al* (2012). 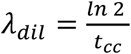 is the dilution constant, where *t*_*cc*_ = 60 minutes. Proteins that obey the following criteria were omitted from the dataset before fitting the model:

i. Less than four measurements.
ii. Proteins that cannot be distinguished based on the respective peptides identified by MS.
iii. Proteins that were identified using less than two peptides.

### Likelihood-ratio test

Early studies have shown that during exponential growth under standard conditions, the *E. coli* proteome is stable, suggesting that for the vast majority of *E. coli*’s proteome under these conditions, the degradation constant is practically zero (Hogness *et al*, 1955; Koch & Levy, 1955; Mandelstam, 1958; Nath & Koch, 1970; Larrabee *et al*, 1980; Camberg *et al*, 2009). We therefore formulated two nested protein degradation models based on equation **(7)**. The first model states that for a given protein, *λ*_*deg*_ = 0:

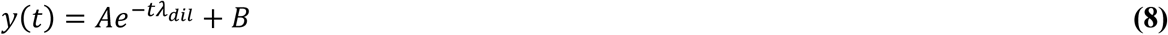

whereas the second model states that *λ*_*deg*_ is a free parameter *λ*_*deg*_ > 0:

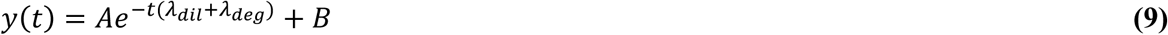

R’s *nls* was used to estimate the parameters fit, using ‘nl2sol’ algorithm from the Port library (Dennis *et al*, 1981). The *nl2sol* algorithm allows setting boundaries for the estimated parameters. For both models, A and B were limited to [0.75, 1.25], [0, 0.4], while for the second model, *λ*_*deg*_ was limited to [0, 100 × *λ*_*dill*_]. These boundaries enabled omitting proteins for which the offset, ***B***, is higher than the initial isotopic ratio, ***A***, as well as to prevent *λ*_*deg*_ from being estimated negative, which is biologically impossible. We constrained *λ*_*deg*_ to be at most 100-fold more effective than *λ*_*dill*_, to prevent the estimation of half-life (see below) to 0, which is also impossible. Using R’s *lrtest* function, likelihood-ratio test was then employed to select the model that best fits the data. The p-values returned by the *lrtest* function were then corrected for multiple testing using Benjamini-Hochberg correction, using R’s *p*.*adjust*. Proteins for which the p-value was equal to, or larger than, 0.05 were labeled as “stable”, whereas the rest of the proteins were labeled as “degradable”. In the case of the degradable group, the fitted *λ*_*deg*_ was used to calculate the half-life time, *t*_1/2_:

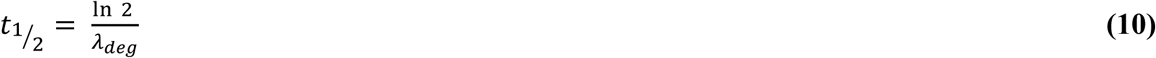

Proteins with fits of low quality (R^2^ < 0.8) in both models were discarded.

### Expectation–maximization algorithm

To determine which degradable proteins are fast- or slow-degrading, we used *mixR* package for expectation–maximization. The calculated *t*_1/2_ was given as an input to *mixR*’s function, *mixfit*, which performs maximum likelihood estimation for various finite mixtures using expectation–maximization algorithm. The statistical significance of the mixture model was estimated using *mixR*’s *bs*.*test* function. A probability threshold of 0.5 was used to attribute each observation to the respective component.

### Enrichment analysis

Gene ontology (GO) molecular function and biological pathways and Kyoto Encyclopedia of Genes and Genomes (KEGG) pathways annotations were analyzed for enrichment using Perseus (Tyanova *et al*, 2016).

### Feature extraction

A total of 188 structural, physical, protein-protein interaction network (PPIN) and physicochemical features were collected (**Supplemental Dataset 2**). Four features describing the intrinsic disorder propensity were extracted using the ESpritz 1.3v webserver (Walsh *et al*, 2011): (1) fraction of disordered amino acids out of the total protein length; (2) total number of disordered segments; (3) total number of disordered segments composed of at least 30 amino acids; (4) total number of disordered segments composed of at least 50 amino acids. Six additional features describing the PPIN of a protein were extracted from the STRING 11.0v database (Szklarczyk *et al*, 2014): (1) the total number of interacting partners of a protein by counting all its neighbors (node connectivity); (2) the average pI of interacting partners; (3) the average molecular weight of interacting partners; (4) the average sequence length of interacting partners; (5) the average disorder among interacting partners was calculated by dividing the total number of disordered amino acids across all interacting partners by the total length of interacting partners; (6) a binary feature describing whether a protein is an isolated node (i.e., a node without neighbors) in the PPIN. Additional 128 PPIN features were extracted using node2vec (Grover & Leskovec, 2016); for these features, isolated nodes were assigned with zeroes. All PPIN-related features were calculated based on the PPIN that was predicted by STRING using only those proteins that were detectable in at least three time points (1,223 proteins). Ten additional features were extracted using ProteinAnalysis class of the Biopython package (Cock *et al*, 2009): (1) molecular weight; (2) average protein aromaticity (Lobry & Gautier, 1994); (3) average protein instability (Guruprasad *et al*, 1990); (4) isoelectric point; (5) average gravy score (Kyte & Doolittle, 1982); (6) average flexibility (Vihinen *et al*, 1994); (7) sequence length; (8) fraction of helix positions; (9) fraction of turn positions; (10) fraction of beta sheet positions. Ten additional features were calculated by dividing each of the Biopython features by the number of interacting partners of each protein. To handle isolated nodes (four proteins), we artificially added one neighbor to all proteins in the network. Twenty additional features are the number of occurrences of each of the 20 amino acids at the second position of the N-terminus. Five additional features are the number of occurrences of each of the 20 amino acids grouped into five physicochemical groups at the second position of the N-terminus: (1) aliphatic (IVL); (2) aromatic (FYWH); (3) charged (KRDE); (4) tiny (GACS) and (5) diverse (TMQNP). Five additional features are the number of occurrences of five different previously described degradation signals: three N-terminal signals termed NM1 (polar-T/ϕ-ϕ-basic-ϕ), NM2 (NH_2_-Met-basic-ϕ-ϕ-ϕ-X5-ϕ), NM3 (ϕ-X-polar-X-polar-X-basic-polar) and two C-terminal signals termed CM1 (LAA-COOH) and CM2 (RRKKAI-COOH).

### Comparative analysis of protein features

One-way ANOVA followed by Tukey’s test was used to test for statistical significance in isoelectric point, mass, percent disorder and number of interacting partners in the PPIN among the three stability groups. Chi square test was used to analyze differences among groups for binary features, e.g., presence/absence of a sequence-related motif. All p values are reported after a Benjamini-Hochberg FDR correction.

### Machine-learning protocol

**C**lassification between several grouping of the proteins were tested: (1) fast versus slow degrading proteins; (2) fast degrading versus stable proteins; (3) slow degrading versus stable proteins; (4) fast degrading versus the rest of the proteins; (5) slow degrading versus the rest of the proteins; (6) stable versus the rest of the proteins; (7) fast degrading versus slow degrading versus stable proteins. We aimed to test whether machine-learning can be used to classify the ORFs into distinct stability groups. We used least absolute shrinkage and selection operator (LASSO) regularized logistic regression (Cox, 1958) for each classification task for its speed, robustness and interpretability. Model training was performed via the Python package ‘scikit-learn’ (Pedregosa *et al*, 2011) using the optimization algorithm ‘liblinear’. The default penalty parameter for regularization was used to train the final model. All learning was based on the 1,149 open reading frames (ORFs) for which we could determine protein degradation-rates (see **Results**).

The performance of the classification was measured in terms of AUC. The performance on the actual data was estimated by 10 repetitions of 10-fold cross-validation, i.e., 90% of the data were randomly chosen for training the model, and the remaining 10% were used for testing the performance of the classification. This was done in a stratified manner, i.e. keeping the relative frequency of the two groups the same in each fold. In each repetition, 10-fold cross-validation is repeated with different randomization of the split to train and test sets. The AUC of each 10-fold cross-validation was calculated by averaging the AUC over the 10 folds. For classification of the three stability groups, the same approach was taken except that scikit-learn’s multinomial logistic regression was used, and the performance of the classification was measured in terms of one-versus-rest AUC, in which the AUC of each class is calculated against the rest and then averaged over the number of classes.

To test whether the AUC is significantly higher than random, class labels (stable/slow degrading/fast degrading) were randomly shuffled among all proteins. The same inference described above was conducted on the permuted data. This was repeated one hundred times. One-way ANOVA followed by Tukey’s test was used to compare the performance of the classifier on the actual versus permuted data.

We tried alternative machine-learning classifications (Random Forest, K nearest neighbors, SVM with various kernels, linear discriminate analysis, and Naïve-Bayes, with and without dimensionality reduction using principle component analysis), which did not provide any significant increase in classification accuracy (not shown). In addition, we considered including various features such as all pairs of amino-acids (400 features) and all triplets (8,000 features). Their inclusion did not contribute to classification accuracy and are hence not shown.

## Results

### Quantification of protein half-lives in *E. coli*

We measured protein half-lives in exponentially growing *E. coli* cells by applying-pulsed SILAC to dividing cells followed by quantitative MS analysis of whole cell extracts as a function of time (see Methods). This is an adaptation to bacteria of the experimental design of Boisvert *et al* (2012). Briefly, the cultures were grown and passaged in the presence of either light (L) or medium (M) lysine isotopes until full incorporation of the label. When the cultures reached mid-exponential phase, the medium of the culture growing with the M lysine was replaced with medium containing the heavier (H) lysine isotope. The rate of protein degradation was inferred from the decreasing ratio of M/L isotopes over time (**Fig. 1**). In total, we identified and quantified 1,602 proteins (**Supplemental Dataset 1**). This value is within the range that was reported for other SILAC (although with only a double labeling, not triple) experiments in bacteria (Soufi *et al*, 2010; Michalik *et al*, 2012). Out of this subset, we estimated the half-life of 1,149 proteins (**Supplemental Dataset 2**, see Methods for filtering criteria).

### Statistical modelling of protein stability reveals that only a small subset of proteins undergoes rapid degradation

The half-life values of the proteins vary dramatically, ranging from minutes to a few days (**Fig. 2A)**. We classified the quantified proteins to either stable or degradable group by selecting one of two nested models of protein degradation (see Methods). The first model states that for a given protein, the exponential decrease in M/L ratio over time is solely governed by protein dilution due to cell division, whereas the second model states that this decrease results from the combined effects of protein dilution and degradation. A total of 408 proteins for which the dilution model was significantly less likely than the degradation and dilution model were termed as degradable, whereas the other 741 proteins were termed stable. This distribution indicates that for the majority of *E. coli* proteins expressed under standard conditions, degradation is undetectable. While most proteins are not degraded under standard conditions, we observed a fraction of unstable proteins that agrees with 2-7% (out of the total protein content) unstable proteins predicted from previous experiments (Nath & Koch, 1970; Larrabee *et al*, 1980; Mosteller *et al*, 1980).

**FIG. 2.**
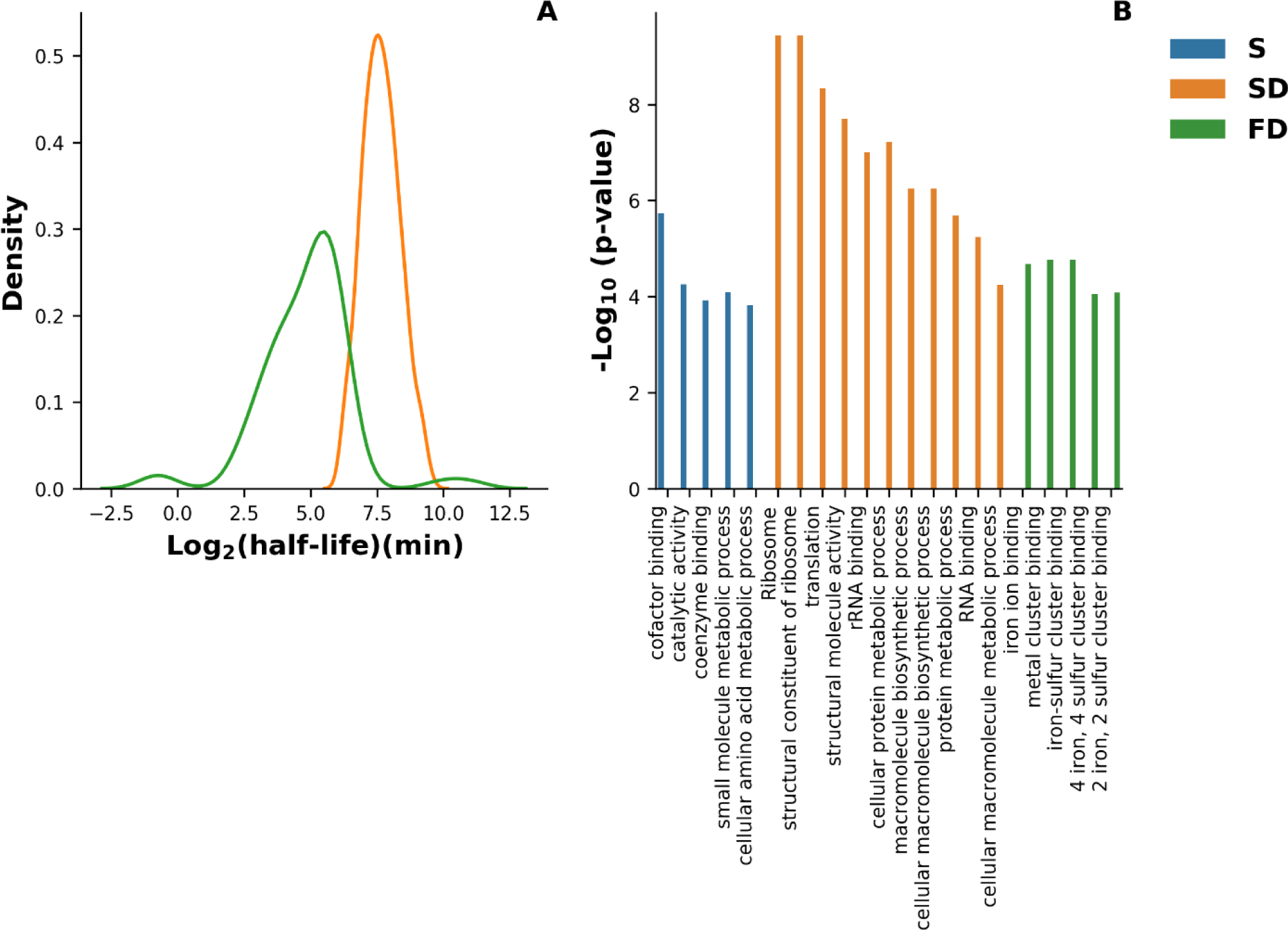
Determination of protein half-lives. **(A)** The distribution of the half-lives of 408 degradable *E. coli* proteins is composed from two distinct subpopulations of slow (SD) (N = 335) and fast (FD) (N = 74) degrading proteins. The bins are log2 increments. Two proteins that had half-lives longer than 16 hours were assigned to the fast degrading proteins by the expectation– maximization algorithm. **(B)** Enrichment analysis demonstrate functional differences between fast degrading and slow degrading/stable proteins based on GO annotations of molecular functions and biological processes as well as KEGG pathways annotations.

Among the fast degrading proteins, we identified several proteins previously reported to have short half-life values, such as RNA polymerase sigma factor (RpoS) and DNA protection during starvation protein (Dps) with half-life of approximately 2 and 10 minutes during exponential phase, respectively (Becker *et al*, 1999; Stephani *et al*, 2003). In this study they were classified as fast degrading proteins with half-life values: 5.8 and 6.5 minutes. An extensive literature survey revealed that out of 72 proteins identified by us as fast-degrading proteins (see below), 21 were previously reported as being prone to degradation, while the remaining 51 are newly identified fast-degrading proteins (**Supplemental Dataset 3**).

It was previously suggested that the degradable fraction of the proteome of *E. coli* is composed of a rapidly and slowly decaying components (Nath & Koch, 1970; Larrabee *et al*, 1980). The alternative hypothesis is that there exists a single component with high variance. We used an expectation–maximization algorithm to estimate the maximum likelihood of the two-component mixture model and compared it against a single component model (see Methods). The expectation–maximization algorithm identified two distinct distributions, ∼Norm(7.64, 0.72) and ∼Norm(5.58, 2) with a log-likelihood of −669.5. Likelihood ratio test by parametric bootstrapping between one-component versus two-component mixture model confirmed the latter (p-value < 0.001), indicating that the degradable group is most likely composed of two distinct protein subpopulations. The expectation–maximization algorithm also assigns probabilities for being a member of a specific distribution. By applying a probability threshold of 0.5, we obtained 334 an4d 74 proteins that are distributed according to ∼Norm(7.64, 0.72) and ∼Norm(5.58, 2), and are therefore termed slow- and fast-degrading, respectively. The proteins YgcE and HolC, which were assigned by the expectation–maximization algorithm to the fast-degrading group, had much longer half-lives than proteins in the slow-degrading group (more than 16 hours). We suspect that these two proteins, which constitute the right-tail density of the half-life distribution of the fast-degrading proteins, were attributed to this group because the extremity of their half-lives is significantly inconsistent with the narrow distribution of half-lives of the slow degrading proteins. We therefore decided to include YgcE and HolC in the group of stable proteins, and thus 72 proteins were classified as fast-degrading proteins.

Since the culture was sampled several times during exponential growth, we hypothesized that most of the stable and slow degrading proteins would be directly involved in growth. It seems unlikely that proteins that are indispensable for growth would be targeted for degradation under conditions in which they are needed most. To test this hypothesis, we analyzed the enrichment of GO molecular function and biological process annotations of stable, slow and fast degrading proteins. Slow and stable proteins were found to be mostly enriched for annotations related to metabolism, biosynthesis and growth, including catalytic activity, cofactor and coenzyme binding, and translation. By contrast, fast degrading proteins were found to be enriched for annotations related to metal binding (**Fig. 2B)**. We suspect that this result reflects the lack of trace metals in the growth medium, suggesting that metal-binding proteins are rapidly degraded in the absence of metals. Such proteins were previously shown to be degraded by AAA+ ATP-dependent proteases (Pruteanu & Baker, 2009). To better understand the biological roles of fast degrading proteins, we also analyzed the annotations that were not significantly enriched. Several fast degrading proteins were found to be either poorly characterized or involved in various processes, including response to diverse stress conditions, including cold, oxidative stress and DNA damage, as well as in proteolysis, regulation of transcription, and biofilm formation. (**Supplemental Dataset 3**).

### Statistical comparison between fast degrading, slow degrading, and stable proteins

Previous studies have reported that structural, physical, and sequence properties, as well as PPIN-associated features, correlate with protein degradation (Dice *et al*, 1979; Miller *et al*, 1987; Varshavsky, 1997; Flynn *et al*, 2003; van der Lee *et al*, 2014; Martin-Perez & Villén, 2017). To test if the fast, slow and stable proteins differ in such properties, we conducted a comparative analysis of various protein-related features across the three protein stability groups. We first analyzed the physicochemical, structural and PPIN properties of the three groups. Stable proteins were found to be slightly more acidic and larger than slow degrading ones (**Fig. 3A and 3B**). However, no significant difference was found between the isoelectric point and mass of fast-degrading and slow-degrading proteins, nor between fast-degrading and stable proteins. This suggests that the degradation of fast degrading proteins is governed by factors other than simple physical properties. Interestingly, stable proteins were found to be significantly less disordered than fast and slow degrading proteins (**Fig. 3C)**. Slow degrading proteins were found to be significantly more connected in the PPIN than fast degrading and stable ones (**Fig. 3D**), suggesting that slow degrading proteins interact with a larger number of proteins, either physically, functionally, or both.

**FIG. 3.**
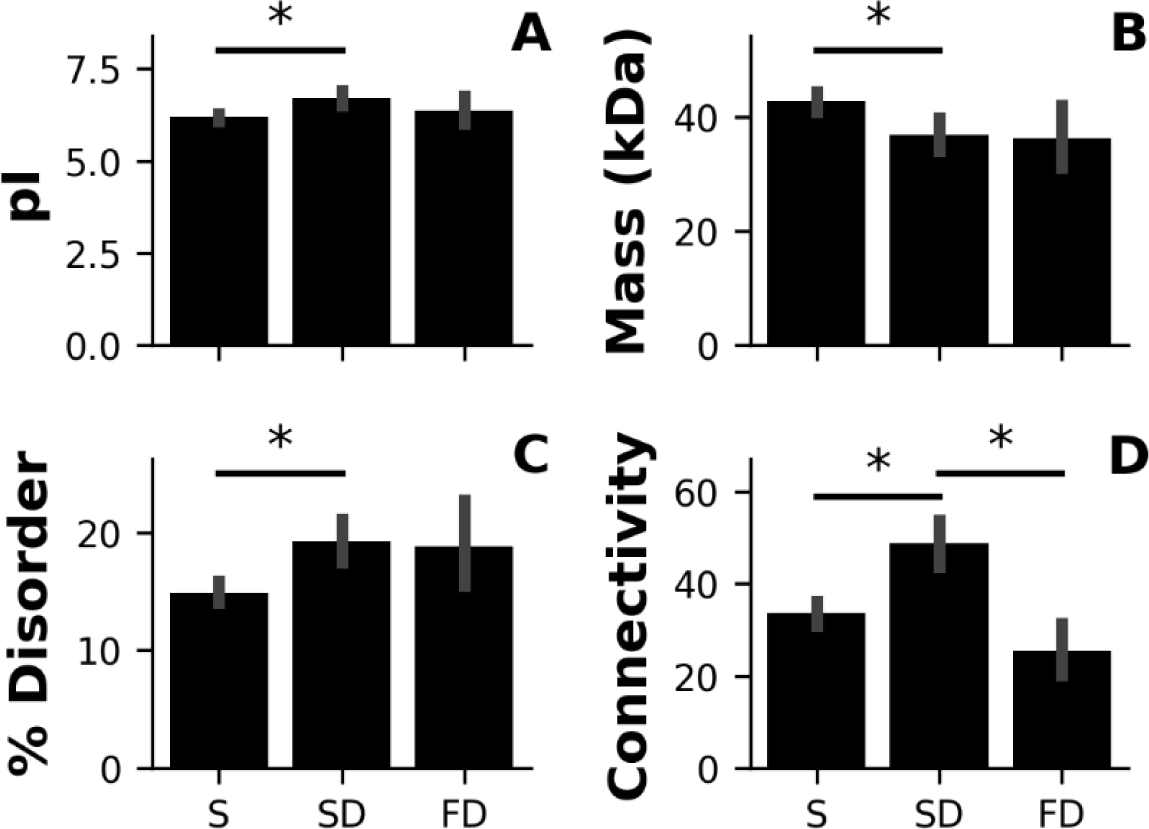
Comparison of protein properties. (A) Isoelectric point, (B) molecular mass, (C) node connectivity and (D) predicted percentage of disordered amino acids. The three stability groups: stable (S), slow degrading (SD) and fast degrading (FD) proteins. * - p-value < 0.005, one-way ANOVA followed by Tukey’s test.

We next analyzed several sequence properties of the three stability groups. The recognition of proteolytic substrates in bacteria is thought to be mediated by short sequence motifs, termed degrons, which are present at the terminal regions of the substrate. Properties that directly or indirectly capture this information are therefore expected to be highly predictive of protein degradation in bacteria. The N-end rule (Varshavsky, 1997) and several other C- and N-termini motifs that were previously reported as important in protein degradation were collected. The frequency of each amino acid at the second position of the N-terminus (after the f-Methionine) was used to capture the N-end rule (see Methods). In addition, the number of occurrences of each amino acid grouped into five physicochemical properties at the second position of the N-terminus was also used to capture the N-end rule. Besides the N-end rule, the number of occurrences of few N- and C-termini sequence motifs that are thought to be recognized by the ClpXP protease were also analyzed (Flynn *et al*, 2003). Together, the N-end rule and ClpXP recognition signals constitute the most established determinants of protein degradation in bacteria. Interestingly, no significant dependency was found between any of these features and protein stability (**Fig. S1**), suggesting that these signals may promote degradation of a small fraction of bacterial proteins.

### Machine-learning to predict fast-degrading proteins

We observed small yet statistically significant differences in the percent of structural disorder, mass, isoelectric point and node connectivity among the various stability groups. We next tested whether these and other presumably informative features could be used to predict the stability category of each protein. To this end, we applied machine learning classification algorithms to find a function between the set of features and the stability group, i.e. to train a machine-learning classifier. An accurate classifier would predict the correct label for “unseen” data. To test the accuracy, a part (fold) of the data is treated as unknown while the remaining folds are used to train the classifier (see Methods). We included all features that are potentially related to protein stability, including physicochemical, structural, sequence and PPIN-related features, as well as features that integrate the node connectivity of each protein with its structural and physicochemical attributes. Overall, 188 features were collected for the classification (**Supplemental Dataset 2**). The performance of the classifier is measured in terms of AUC, where an AUC of 1 indicates perfect classification and an AUC of 0.5 corresponds to a random classification. The highest accuracies were obtained when fast-degrading proteins were not grouped with either slow-degrading/stable ones. The highest score (AUC 0.74 ± 0.01) was obtained when comparing fast versus slow-degrading proteins (**Fig. 4A**). All these comparisons are significantly better than a classifier trained on permuted data sets (all p-values < 0.001), conforming that the feature set used for training the classifier contains features that are significantly correlated with protein degradation.. The quality of discrimination between the fast degrading and the slow degrading/stable proteins is of special interest, because good discrimination will enable the computational prediction of fast-degrading proteins. In this setting, our classifier achieved 0.72 ± 0.01 AUC, suggesting that intrinsic protein properties as well as PPIN-related features are predictive of protein stability in *E. coli*. Interestingly, the most informative features that discriminate fast degrading proteins from slow degrading and stable ones were PPIN-related features, suggesting that fast-degrading proteins share similar network properties (**Fig. 4B**).

**FIG. 4.**
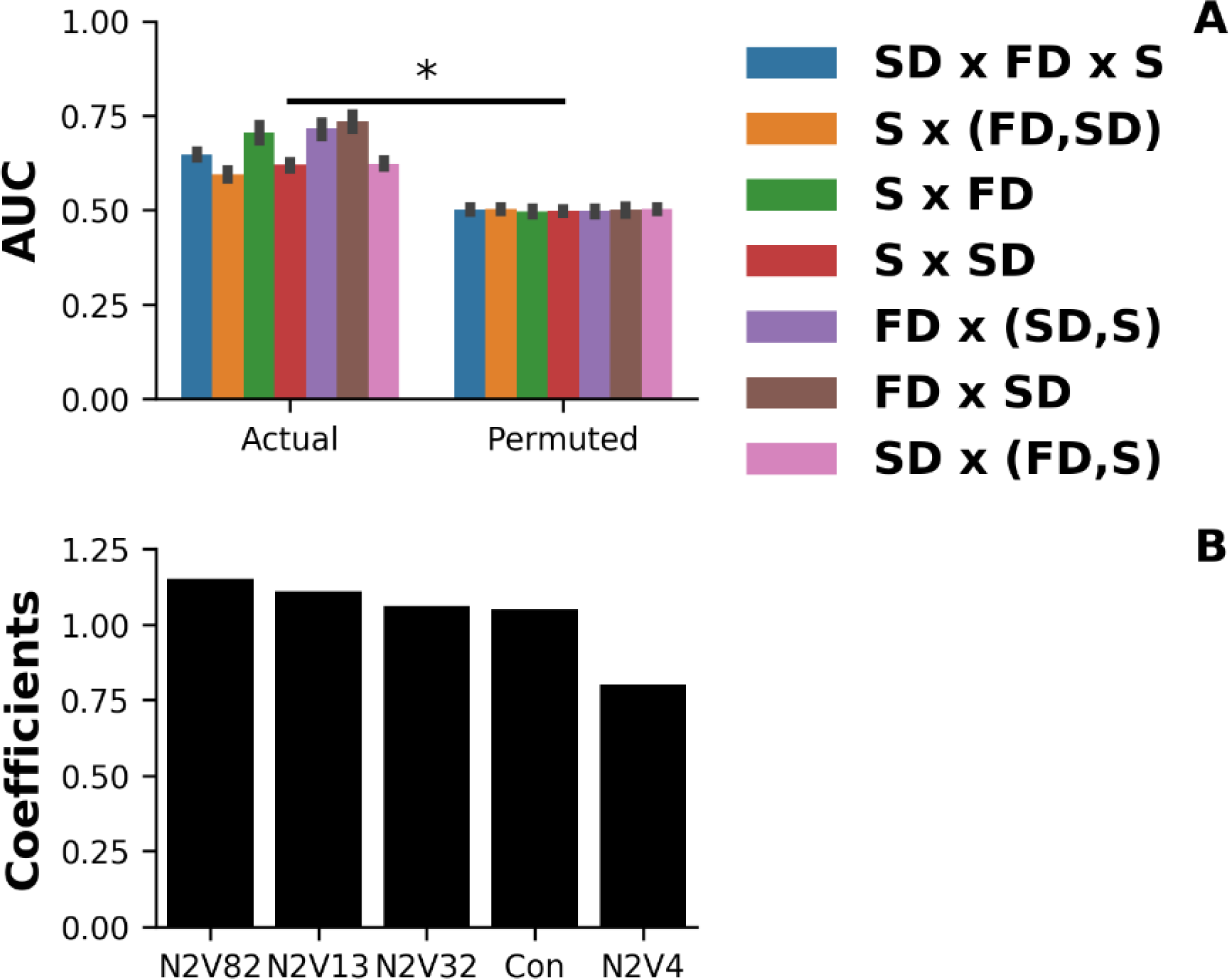
PPIN and physical protein features discriminate fast degrading proteins from stable/slow degrading ones. **(A)** Classification of proteins to stable (S), slow degrading (SD) and fast degrading (FD) using logistic regression trained with 188 physicochemical, structural and PPIN-related features is significantly better than random. Stability groups (S, SD, FD) are in round brackets when one stability group was compared against the rest of the groups. All models trained with the actual data were compared using paired t-test to their corresponding permuted dataset. * - p-value <0.0001, paired t-test followed by FDR correction. The performances obtained using the actual datasets were significantly higher than their corresponding permuted datasets. The AUC of the actual data was estimated by ten repeats of 10-fold cross-validation while the AUC corresponding permuted data is an average of one hundred repeats of 10-fold cross-validation, where each repeat is a different permutation of the class labels. **(B)** Top five features associated with the fast degrading group versus the rest. Con – node connectivity in the PPIN. Node2vec (N2V) features-#82 (N2V82), #13 (N2V13), #32 (N2V32) and #4 (N2V4).

## Discussion

The degradation of intracellular proteins is important for the regulation of cellular processes and serves as a mechanism for protein quality control. Hence, the quantification of protein half-lives and the elucidation of factors determining degradation dynamics are critical for the understanding of protein activity regulation. In this work, pulsed SILAC followed by LC-MS were applied to explore protein degradation in *E. coli* during the exponential phase of growth. This enabled monitoring the degradation of proteins that were present in the cell at early-mid exponential phase of growth, at the proteome level. A key step for understanding protein degradation is the reliable quantification of the half-lives of all proteins expressed under a given condition. To achieve this, we determined and modelled the degradation of 1,149 proteins, which is nearly half of the expressed proteins in *E. coli* (Li *et al*, 2014), providing the largest dataset of its kind for protein half-lives for this species. The use of log-likelihood ratio test combined with expectation–maximization to choose the most likely mode of degradation for each protein revealed three distinct stability groups: stable, slow degrading and fast degrading proteins. The vast majority of the proteins were classified as highly stable or slow degrading (66% and 29.1%, respectively). The remaining 6.3% were found to be fast degrading, with half-lives ranging from 70 minutes to less than a minute (**Fig. 2A**). These values are in agreement with an early study that found that only 2-7% of the *E. coli* protein content undergoes rapid degradation during the exponential growth phase (Nath & Koch, 1970). We assume that most of the proteins that were not identified in this study are either not expressed or are too unstable to be detected in our experimental setting. In this context, we encountered what seems to be a typical limitation of pulsed SILAC methods (Boisvert *et al*, 2012; Michalik *et al*, 2012), in which respective peptides are undetectable for certain proteins in some of the sampled time points, leading to some loss of information.

A similar pulsed SILAC approach was previously taken to track protein degradation at the transition from exponential to stationary phase of growth in *Staphylococcus aureus* (Michalik *et al*, 2012). In this setting, most proteins that undergo rapid degradation are proteins that are essential in substantial amounts during the exponential phase, such as ribosomal proteins, and anabolic or catabolic enzymes. In our experimental setting, proteins required for growth were found to be mostly stable or slow degrading, while the fast degrading proteins had diverse roles including metal binding, response to various stresses, and transcriptional regulation (**Fig. 2B and Supplemental Dataset 3**). This suggests that protein degradation is differentially regulated at the various stages of growth, and that proteins that are unstable during growth may become stable under stress or starvation, and vice versa.

The fast-degrading proteins have high turnover during growth. What may be the biological significance of such a phenomenon, i.e. why should evolution favor a state in which proteins are continuously transcribed and translated only to immediately be degraded? We propose six possible explanations: (i) These proteins harbor degrons recognized by the AAA+ ATP-dependent proteases which could not be eliminated in the course of evolution due to structural or functional constraints. (ii) Rapid accumulation of fast degrading proteins can be achieved by stopping their degradation, e.g., by the inhibition of specific proteases or modulation of adaptors. Thus, the degradation of such proteins is used as a regulatory switch that keeps their concentration low at exponential phase, yet allows a rapid increase in their concentration upon an environmental change (Zgurskaya *et al*, 1997; Mandel & Silhavy, 2005). (iii) Proteins that are involved in specific steps of the cell-cycle might oscillate between cycles, which may cause us to identify them as fast-degrading proteins (Camberg *et al*, 2009, 2011). (iv) Protein degradation adjusts the level of proteins which are members in heterocomplexes and are synthesized at different levels. (v) These proteins are prone to misfolding under exponential growth conditions, and most of the proteolysis is of the misfolded variants. (vi) The instability of these proteins is protease-independent. Clearly, the current data do not allow us to determine the relative contribution of each of these possible factors.

Proteolysis was previously suggested to have a role in regulating the activity of RpoS and Dps proteins. RpoS regulates gene cascades that are involved in response to various stress conditions, including oxidative stress, extreme temperature, pH, and osmolarity as well as DNA damage. The Dps protein binds and thereby protects DNA from oxidative stress. It was suggested that inhibition of their constant degradation by AAA+ ATP-dependent proteases during the exponential phase is important for their rapid accumulation following stress, which in turn enables them to respond quickly to the stress signal (Neher *et al*, 2006). We note that testing the biological effect of protein stability and the role of specific residues in governing protein stability, *in-vivo*, is a challenging task since residues may play multiple roles, e.g., in protein folding, interaction with other molecules, and in stability (Becker *et al*, 1999).

Studying protease-independent stability can be conducted by systematic determination of the stability of purified proteins *in-vitro*. Another possibility is to study degradation rates *in-vivo*, in which all AAA+ ATP-dependent proteases are knocked out. In the case of the essential FtsH protease (Baba *et al*, 2006)., such studies can be conducted in conditional mutants for this gene (Fischer *et al*, 2002).The effect of various physical (temperature, osmolarity) and biological (medium composition, introduction of stress) factors on protein stability remains to be studied. Moreover, the effect of ATP-independent proteases remains to be discovered. Finally, it is of interest to discover if, and how, bacteriophages manipulate protein degradation rates to their benefit.

Once we obtained reliable information on protein degradation, we could focus on the more challenging problem of identifying key differences between fast-degrading proteins and the rest of the quantified proteome and using them for prediction. A prerequisite for this challenge is to objectively sort the proteins (in the training set) into different stability groups. Here we employed likelihood ratio tests together with expectation maximization, thereby avoiding arbitrary cutoffs for discriminating between the stability groups. This objective criterion revealed the existence of three distinct stability groups. We collected several features previously reported as correlated with degradation, as well as other potentially predictive ones. We showed that physicochemical and PPIN properties are more correlated with degradation than previously described degrons (**Fig. S1 and 3**). This implies that both substrate specificity and substrate selectivity of AAA+ ATP-dependent proteases are broader than previously thought. Our machine-learning algorithm combines both structural and physicochemical features with PPIN-related features to classify proteins to different stability groups (**Fig. 4**). It would be interesting to estimate how well our machine-learning approach generalizes to evolutionary-related proteobacteria and diverged bacterial species.

## Acknowledgements

Israel Science Foundation (ISF) [802/16 to T.P.]; Edmond J. Safra Center for Bioinformatics at Tel Aviv University Fellowship. The authors declare that they have no conflict of interest.

---

**FIG. S1.**
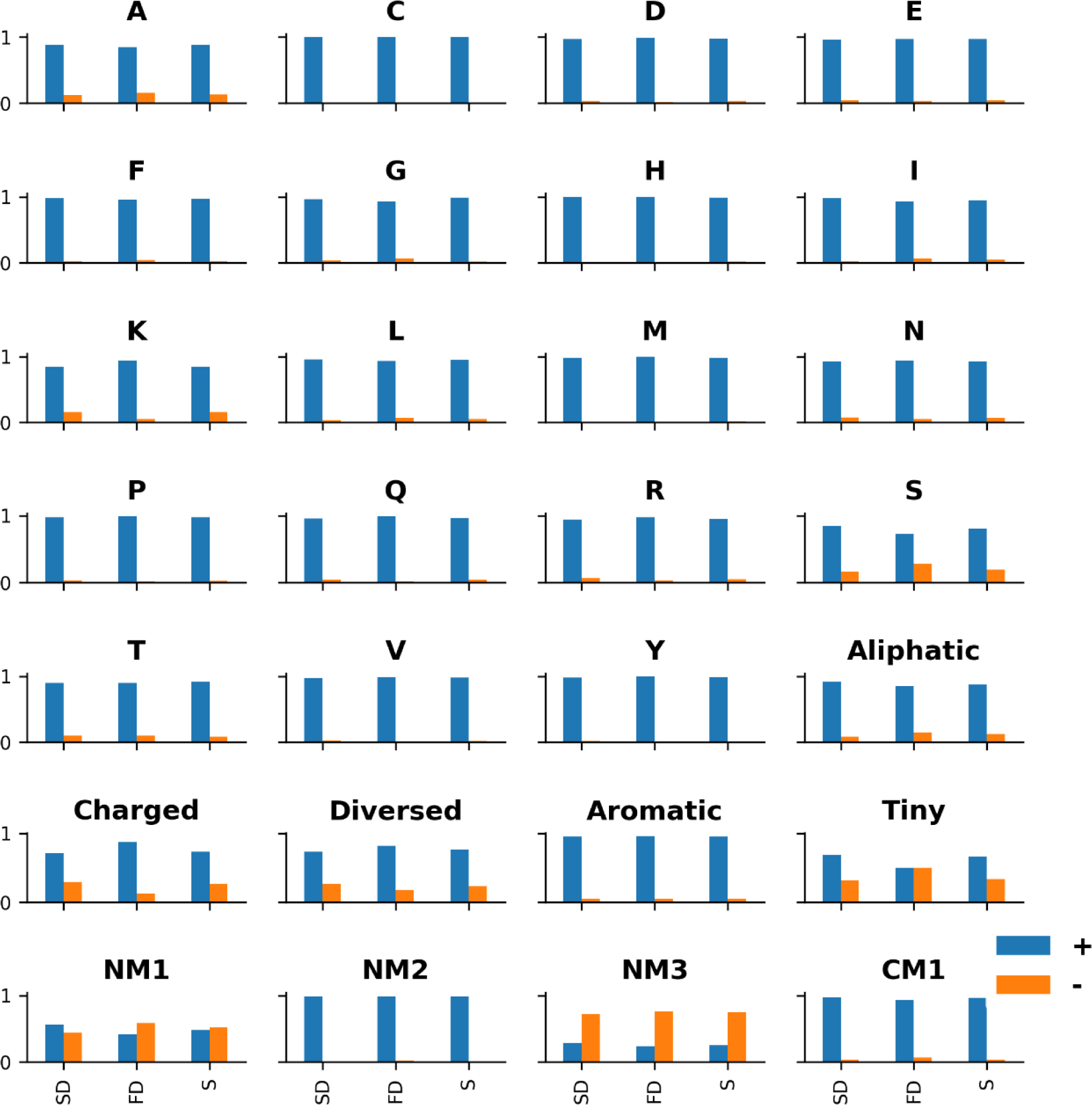
Normalized frequency of previously reported characteristics of unstable proteins in bacteria. The N-end rule (Varshavsky, 1997), three N-termini (NM1-3) and two C-termini (CM1-2) recognition sequences of ClpXP (Flynn *et al*, 2003). The number of occurrences of each of the 20 amino acids, represented by single letters (ACDEFGHIKLMNPQRSTVY) as well as the number of occurrences of Aliphatic (IVL), Aromatic (FYWH), Charged (KRDE), Tiny (GACS) or Diverse (TMQNP) amino acids at the second position of the N-terminus summarize the N-end rule. The amino acids tryptophan (W) was not observed at this position, and CM2 motif was not found, in any of the proteins. Protein stability was not found to be dependent on any of these sequence motifs (Chi square test followed by Benjamini-Hochberg FDR correction). Stable (S), slow degrading (SD) and fast degrading (FD) proteins. The presence or absence of a sequence motif is designated by + and -, respectively.

## Bibliography

Arends J, Griego M, Thomanek N, Lindemann C, Kutscher B, Meyer HE, Narberhaus F (2018) An integrated proteomic approach uncovers novel substrates and functions of the Lon protease in Escherichia coli. Proteomicss 18: 1800080

Baba T, Ara T, Hasegawa M, Takai Y, Okumura Y, Baba M, Datsenko KA, Tomita M, Wanner BL, Mori H (2006) Construction of Escherichia coli K-12 in-frame, single-gene knockout mutants: The Keio collection. Mol Syst Biol 2: 2006–0008

Bachmair A, Finley D, Varshavsky A (1986) In vivo half-life of a protein is a function of its amino-terminal residue. Science 234: 179–186

Baker TA, Sauer RT (2006) ATP-dependent proteases of bacteria: recognition logic and operating principles. Trends Biochem Sci 31: 647–653

Becker G, Klauck E, Hengge-Aronis R (1999) Regulation of RpoS proteolysis in Escherichia coli: The response regulator RssB is a recognition factor that interacts with the turnover element in RpoS. Proc Natl Acad Sci U S A 96: 6439–6444

Boisvert FM, Ahmad Y, Gierliński M, Charrière F, Lamont D, Scott M, Barton G, Lamond AI (2012) A quantitative spatial proteomics analysis of proteome turnover in human cells. Mol Cell Proteomics 11: M111–011429

Burstein D, Gould SB, Zimorski V, Kloesges T, Kiosse F, Major P, Martin WF, Pupko T, Dagan T (2012) A machine learning approach to identify hydrogenosomal proteins in Trichomonas vaginalis. Eukaryot Cell 11: 217–228

Burstein D, Zusman T, Degtyar E, Viner R, Segal G, Pupko T (2009) Genome-scale identification of Legionella pneumophila effectors using a machine learning approach. PLoS Pathog 5: e1000508

Burton RE, Baker TA, Sauer RT (2005) Nucleotide-dependent substrate recognition by the AAA+ HslUV protease. Nat Struct Mol Biol 12: 245–251

Camberg JL, Hoskins JR, Wickner S (2009) ClpXP protease degrades the cytoskeletal protein, FtsZ, and modulates FtsZ polymer dynamics. Proc Natl Acad Sci U S A 106: 10614–10619

Camberg JL, Hoskins JR, Wickner S (2011) The Interplay of ClpXP with the cell division machinery in Escherichia coli. J Bacteriol 193: 1911–1918

Cheng X, Xiao X, Chou KC (2018) pLoc-mGneg: Predict subcellular localization of Gram-negative bacterial proteins by deep gene ontology learning via general PseAAC. Genomics 110: 231–239

Christiano R, Nagaraj N, Fröhlich F, Walther TC (2014) Global proteome turnover analyses of the Yeasts S. cerevisiae and S. pombe. Cell Rep 9: 1959–1965

Cock PJA, Antao T, Chang JT, Chapman BA, Cox CJ, Dalke A, Friedberg I, Hamelryck T, Kauff F, Wilczynski B et al (2009) BioPython: Freely available Python tools for computational molecular biology and bioinformatics. Bioinformatics 25: 1422–1423

Conlon BP, Nakayasu ES, Fleck LE, Lafleur MD, Isabella VM, Coleman K, Leonard SN, Smith RD, Adkins JN, Lewis K (2013) Activated ClpP kills persisters and eradicates a chronic biofilm infection. Nature 503: 365–370

Cox DR (1958) The regression analysis of binary sequences. J R Stat Soc Ser B 20: 215–232

Cox J, Mann M (2008) MaxQuant enables high peptide identification rates, individualized p.p.b.- range mass accuracies and proteome-wide protein quantification. Nat Biotechnol 26: 1367–1372

Cox J, Neuhauser N, Michalski A, Scheltema RA, Olsen J V, Mann M (2011) Andromeda: A peptide search engine integrated into the MaxQuant environment. J Proteome Res 10: 1794–1805

Dennis JE, Gay DM, Welsch RE (1981) Algorithm 573: NL2SOL—An Adaptive Nonlinear Least-Squares Algorithm [E4]. ACM Trans Math Softw 7: 369–383

Dice JF, Hess EJ, Goldberg AL (1979) Studies on the relationship between the degradative rates of proteins in vivo and their isoelectric points. Biochem J 178: 305–312

Flynn JM, Levchenko I, Seidel M, Wickner SH, Sauer RT, Baker TA (2001) Overlapping recognition determinants within the ssrA degradation tag allow modulation of proteolysis. Proc Natl Acad Sci U S A 98: 10584–10589

Flynn JM, Neher SB, Kim YI, Sauer RT, Baker TA (2003) Proteomic discovery of cellular substrates of the ClpXP protease reveals five classes of ClpX-recognition signals. Mol Cell 11: 671–683

Goldberg AL (2003) Protein degradation and protection against misfolded or damaged proteins. Nature 426: 895–899

Gsponer J, Futschik ME, Teichmann SA, Babu MM (2008) Tight regulation of unstructured proteins: From transcript synthesis to protein degradation. Science 322: 1365–1368

Gur E, Biran D, Ron EZ (2011) Regulated proteolysis in Gram-negative bacteria-how and when? Nat Rev Microbiol 9: 839–848

Gur E, Sauer RT (2008) Recognition of misfolded proteins by Lon, a AAA+ protease. Genes Dev 22: 2267–2277

Hayes WS, Borodovsky M (1998) How to interpret an anonymous bacterial genome: Machine learning approach to gene identification. Genome Res 8: 1154–1171

Hershko A (1991) The ubiquitin pathway for protein degradation. Trends Biochem Sci 16: 265–268

Hogness DS, Cohn M, Monod J (1955) Studies on the induced synthesis of β-galactosidase in Escherichia coli: The kinetics and mechanism of sulfur incorporation. BBA - Biochim Biophys Acta 16: 99–116

Hoskins JR, Kim SY, Wickner S (2000) Substrate recognition by the ClpA chaperone component of ClpAP protease. J Biol Chem 275: 35361–35367

Hosoi T, Ozawa K (2010) Endoplasmic reticulum stress in disease: mechanisms and therapeutic opportunities. Clin Sci 118:19–29

Huang DW, Sherman BT, Lempicki RA (2009) Systematic and integrative analysis of large gene lists using DAVID bioinformatics resources. Nat Protoc 4: 44–57

Ishii Y, Sonezaki S, Iwasaki Y, Miyata Y, Akita K, Kato Y, Amano F (2000) Regulatory role of C-terminal residues of SulA in its degradation by Lon protease in Escherichia coli. J Biochem 127: 837–844

Jones E, Oliphant T, Peterson P (2001) SciPy: Open source scientific tools for Python. https://www.scipy.org

Jovanovic M, Rooney MS, Mertins P, Przybylski D, Chevrier N, Satija R, Rodriguez EH, Fields AP, Schwartz S, Raychowdhury R (2015) Dynamic profiling of the protein life cycle in response to pathogens. Science 347: 1112

Keiler KC, Waller PRH, Sauer RT (1996) Role of a peptide tagging system in degradation of proteins synthesized from damaged messenger RNA. Science 271: 990–993

Koch AL, Levy HR (1955) Protein turnover in growing cultures of Escherichia coli. J Biol Chem 217: 947–957

Koren I, Timms RT, Kula T, Xu Q, Li MZ, Elledge SJ (2018) The Eukaryotic Proteome Is Shaped by E3 Ubiquitin Ligases Targeting C-Terminal Degrons. Cell 173: 1622-1635.e14

Kyte J, Doolittle RF (1982) A simple method for displaying the hydropathic character of a protein. J Mol Biol 157: 105–132

Larrabee KL, Phillips JO, Williams GJ, Larrabee AR (1980) The relative rates of protein synthesis and degradation in a growing culture of Escherichia coli. J Biol Chem 255: 4125–4130

van der Lee R, Lang B, Kruse K, Gsponer J, de Groot NS, Huynen MA, Matouschek A, Fuxreiter M, Babu MM (2014) Intrinsically disordered segments affect protein half-life in the cell and during evolution. Cell Rep 8: 1832–1844

Li GW, Burkhardt D, Gross C, Weissman JS (2014) Quantifying absolute protein synthesis rates reveals principles underlying allocation of cellular resources. Cell 157: 624–635

Lin HC, Yeh CW, Chen YF, Lee TT, Hsieh PY, Rusnac D V., Lin SY, Elledge SJ, Zheng N, Yen HCS (2018) C-Terminal end-directed protein elimination by CRL2 ubiquitin ligases. Mol Cell 70: 602-613.e3

Mahmoud SA, Chien P (2018) Regulated proteolysis in bacteria. Annu Rev Biochem 87: 677–696

Mandel MJ, Silhavy TJ (2005) Starvation for different nutrients in Escherichia coli results in differential modulation of RpoS levels and stability. J Bacteriol 187: 434–442

Mandelstam J (1958) Turnover of protein in growing and non-growing populations of Escherichia coli. Biochem J 69: 110–119

Mann M (2006) Functional and quantitative proteomics using SILAC. Nat Rev Mol Cell Biol 7: 952–958

Martin-Perez M, Villén J (2017) Determinants and regulation of protein turnover in yeast. Cell Syst 5: 283–294

Mathieson T, Franken H, Kosinski J, Kurzawa N, Zinn N, Sweetman G, Poeckel D, Ratnu VS, Schramm M, Becher I et al (2018) Systematic analysis of protein turnover in primary cells. Nat Commun 9: 689

Maupin-Furlow J (2012) Proteasomes and protein conjugation across domains of life. Nat Rev Microbiol 10: 100

Michalik S, Bernhardt J, Otto A, Moche M, Becher D, Meyer H, Lalk M, Schurmann C, Schlüter R, Kock H et al (2012) Life and death of proteins: A case study of glucose-starved Staphylococcus aureus. Mol Cell Proteomics 11: 558–570

Miller ML, Soufi B, Jers C, Blom N, Macek B, Mijakovic I (2009) NetPhosBac - A predictor for Ser/Thr phosphorylation sites in bacterial proteins. Proteomics 9: 116–125

Miller S, Lesk AM, Janin J, Chothia C (1987) The accessible surface area and stability of oligomeric proteins. Nature 328: 834

Mosteller RD, Goldstein R V, Nishimoto KR (1980) Metabolism of individual proteins in exponentially growing Escherichia coli. J Biol Chem 255: 2524–2532

Nanni L, Lumini A, Gupta D, Garg A (2012) Identifying bacterial virulent proteins by fusing a set of classifiers based on variants of Chou’s Pseudo amino acid composition and on evolutionary information. IEEE/ACM Trans Comput Biol Bioinforma 9: 467–475

Nath K, Koch AL (1970) Protein degradation in Escherichia coli I. Measurement of rapidly and slowly decaying components. J Biol Chem 245: 2889–2900

Neher SB, Villén J, Oakes EC, Bakalarski CE, Sauer RT, Gygi SP, Baker TA (2006) Proteomic profiling of ClpXP substrates after DNA damage reveals extensive instability within SOS Regulon. Mol Cell 22: 193–204

Pedregosa F, Grisel O, Weiss R, Passos A, Brucher M, Varoquax G, Gramfort A, Michel V, Thirion B, Grisel O et al (2011) Scikit-learn: Machine learning in Python. J Mach Learn Res 12: 2825–2830

Price JC, Guan S, Burlingame A, Prusiner SB, Ghaemmaghami S (2010) Analysis of proteome dynamics in the mouse brain. Proc Natl Acad Sci U S A 107: 14508–14513

Pruteanu M, Baker TA (2009) Proteolysis in the SOS response and metal homeostasis in Escherichia coli. Res Microbiol 160: 677–683

Rogers S, Wells R, Rechsteiner M (1986) Amino acid sequences common to rapidly degraded proteins: The PEST hypothesis. Science 234: 364–368

Rubinsztein DC (2006) The roles of intracellular protein-degradation pathways in neurodegeneration. Nature 443: 780

Schwanhäusser B, Busse D, Li N, Dittmar G, Schuchhardt J, Wolf J, Chen W, Selbach M (2011) Global quantification of mammalian gene expression control. Nature 473: 337

Schwanhäusser B, Gossen M, Dittmar G, Selbach M (2009) Global analysis of cellular protein translation by pulsed SILAC. Proteomics 9: 205–209

Shah IM, Wolf RE (2006) Sequence requirements for Lon-dependent degradation of the Escherichia coli transcription activator SoxS: Identification of the SoxS residues critical to proteolysis and specific inhibition of in vitro degradation by a peptide comprised of the N-terminal. J Mol Biol 357: 718–731

Soufi B, Kumar C, Gnad F, Mann M, Mijakovic I, MacEk B (2010) Stable isotope labeling by amino acids in cell culture (SILAC) applied to quantitative proteomics of Bacillus subtilis. J Proteome Res 9: 3638–3646

Swovick K, Welle KA, Hryhorenko JR, Seluanov A, Gorbunova V, Ghaemmaghami S (2018) Cross-species comparison of proteome turnover kinetics. Mol Cell Proteomics 17: 580–591

Szklarczyk D, Franceschini A, Wyder S, Forslund K, Heller D, Huerta-Cepas J, Simonovic M, Roth A, Santos A, Tsafou KP (2014) STRING v10: protein–protein interaction networks, integrated over the tree of life. Nucleic Acids Res 43: D447–D452

Teper D, Burstein D, Salomon D, Gershovitz M, Pupko T, Sessa G (2016) Identification of novel Xanthomonas euvesicatoria type III effector proteins by a machine-learning approach. Mol Plant Pathol 17: 398–411

Tompa P, Prilusky J, Silman I, Sussman JL (2008) Structural disorder serves as a weak signal for intracellular protein degradation. Proteins Struct Funct Bioinforma 71: 903–909

Tyanova S, Temu T, Sinitcyn P, Carlson A, Hein MY, Geiger T, Mann M, Cox J (2016) The Perseus computational platform for comprehensive analysis of (prote)omics data. Nat Methods 13: 731–740

Varshavsky A (1997) The N-end rule pathway of protein degradation. Genes to Cells 2: 13–28

Walsh I, Martin AJM, Di Domenico T, Tosatto SCE (2011) ESpritz: accurate and fast prediction of protein disorder. Bioinformatics 28: 503–509

Westphal K, Langklotz S, Thomanek N, Narberhaus F (2012) A trapping approach reveals novel substrates and physiological functions of Escherichia coli the essential protease Ftsh in Escherichia coli. J Biol Chem 287: 42962–42971

Zgurskaya HI, Keyhan M, Matin A (1997) The σ(s) level in starving cells increases solely as a result of its increased stability, despite decreased synthesis. Mol Microbiol 24: 643–651

